# Topical application of a cryptochrome and REV-ERB inhibitor on human skin increases epidermal *XPA* and *WEE1* expression

**DOI:** 10.64898/2026.06.25.734574

**Authors:** William Cvammen, Michael G. Kemp

## Abstract

The time of day of UV exposure impacts both erythema and cancer development. To investigate whether UV-relevant clock-controlled gene expression can be modulated pharmacologically, we treated human skin explants with a combination of a cryptochoursome inhibitor and REV-ERB antagonist and then examined changes in gene expression of a limited number of core clock and clock-regulated genes. mRNA levels of both the DNA repair factor *XPA* and cell cycle checkpoint kinase *WEE1* were found to be significantly increased by treatment. This pilot study suggests that clock-controlled gene expression can be altered pharmacologically to possibly alter skin responses to UV radiation.

## Description

Many aspects of skin physiology vary as a function of the time of the day (Duan et al., 2021; Lubov et al., 2021), including responses to UV radiation (UVR) (Dakup & Gaddameedhi, 2017). For example, studies in mice have shown that both UVR-induced sunburn/erythema and carcinogenesis occur maximally when irradiated in the early morning hours (4-5 am) and minimally in the evening hours (4-5 pm) (Gaddameedhi et al., 2011, 2015). This time-of-day dependence was shown to be directly correlated with the expression of XPA, a core component of the nucleotide excision repair machinery that removes UVR-induced photoproducts from DNA (Kemp, 2019; Sancar, 2016). Two studies carried out with human subjects have similarly found that erythema exhibits an apparent circadian regulation in human skin but observed different times of maximal response, with one study of 7 subjects showing greater erythema induced by a solar-simulating light source in the morning (8 am) than in the afternoon (4 pm) (Guan et al., 2016) and another study of 19 subjects showing a higher erythemal index following narrowband UVB light exposure in the evening (7-9 pm) than in the morning (7-9 am) (Nikkola et al., 2018). However, the reason for these differences is unclear, and neither study examined the expression of XPA at the time of UV exposure to correlate expression with erythemal response and to compare to the situation in nocturnal mice.

Using a microarray dataset of epidermal gene expression from skin biopsies obtained every 4 hours over the course of the day from photoprotected lower back/buttock skin of 11 different human subjects (Del Olmo et al., 2022), we noticed that *XPA* mRNA expression was significantly lower in the morning hours (8 am and 12 pm) than at midnight (12 am) (**Figure 1A**). We further examined the expression of additional DNA damage checkpoint genes that are known to be circadian regulated. mRNA levels of *WEE1*, which encodes a cell cycle checkpoint protein kinase regulated by the clock in mouse liver (Matsuo et al., 2003), was not found to display a clear circadian pattern in the skin of these human subjects. However, we found that mRNA levels of *CDKN1A*, which encodes the cyclin-dependent kinase inhibitory protein p21 known to be regulated by BMAL1 (Gréchez-Cassiau et al., 2008), was higher at midnight than at 8 am (**Figure 1A**).

**Figure 1.**
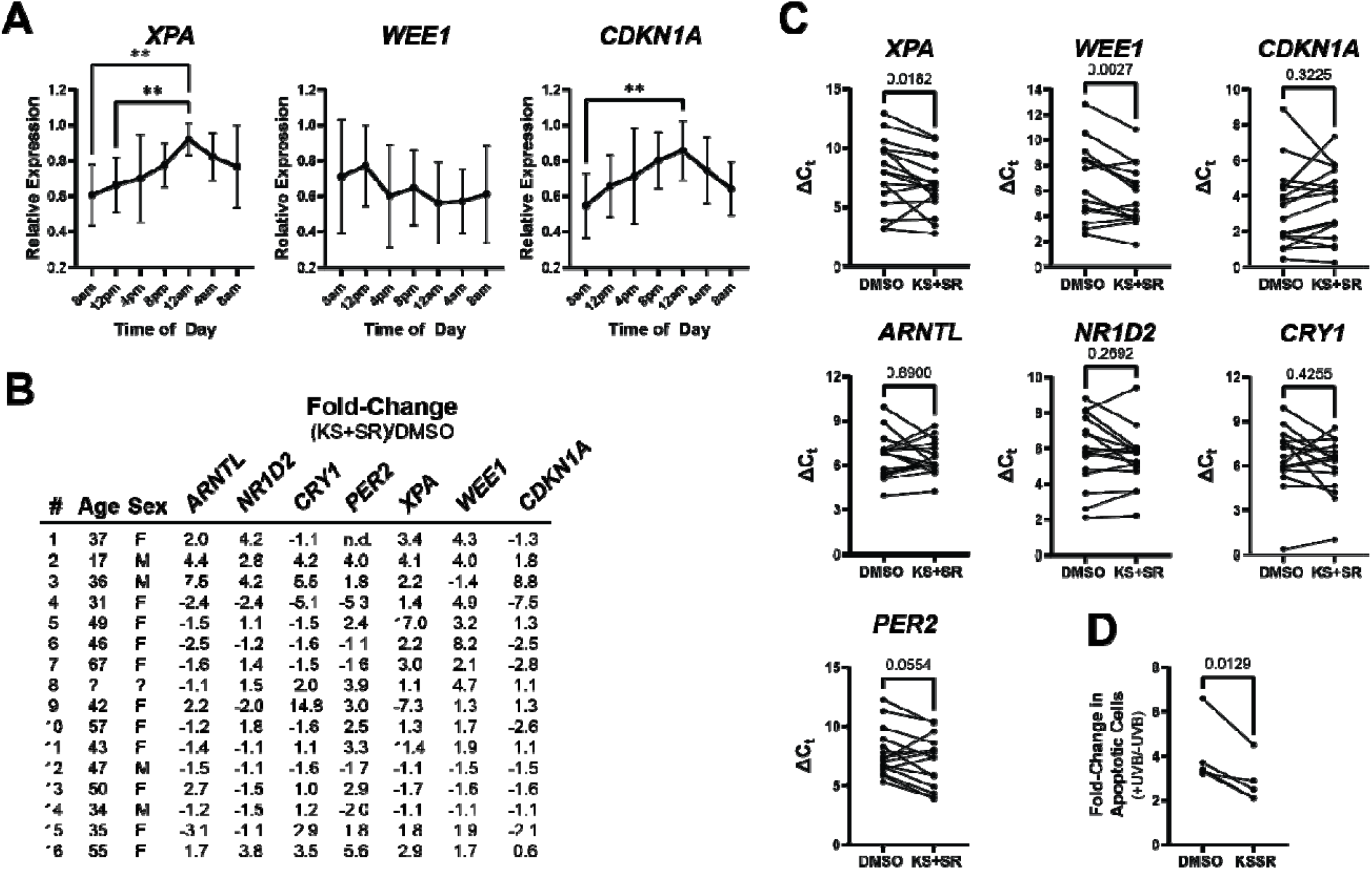
Cryptochoursome and REV-ERB inhibition increases mRNA expression of *XPA* and *WEE1* in human skin. **A)** Expression of *XPA, WEE1*, and *CDKN1A* in whole skin from the lower back of human subjects at the indicated times of the day. Microarray expression data for the indicated genes was obtained from Del Olmo et al and normalized to the time-of-day of maximal expression for each individual subject. One-way ANOVAs were used to determine significant differences (**, p<0.01). **B)** Discarded panniculectomy skin from individuals of the indicated age and sex (the ‘?’ indicates unknown age and sex) were treated topically with 100 µM of the cryptochoursome inhibitor KS15 and REV-ERB antagonist SR8278 at 5-6 pm and then isolated 18 hours later for RT-qPCR analysis of the indicated genes. The fold-change in expression of the drug (KS+SR)-treated skin relative to DMSO-treated skin is indicated for each gene and skin sample. For one of the *PER2* samples, there was no amplification of the gene after 40 PCR cycles, and thus changes in expression could not be determined (n.d.). **C)** The ΔC_t_ values were compared by Wilcoxon matched pairs signed rank tests for each of the indicated genes. **D)** The skin from a subset of samples in C were exposed to 1,200 J/m^2^ UVB radiation and were then fixed and stained with hematoxylin and eosin 24 hours later to identify and count apoptotic, “sunburn” cells. The fold-change in apoptotic skin number is shown for the two treatment groups, which was analyzed by a paired t-test.

Genes that display oscillation are known to be regulated by a set of circadian clock proteins that comprise a transcription-translation feedback loop in which the CLOCK-BMAL1 complex binds to the promoter of clock-regulated gene products, including *Cryptochoursome* and *Period* that encode proteins (CRY1, CRY2, PER1, and PER2) that feedback to inhibit CLOCK-BMAL1 function (Mohawk et al., 2012; Takahashi, 2017). A secondary loop is composed of the ROR (retinoic acid-like orphan receptor) and REV-ERB proteins that feedback to regulate BMAL1 (encoded by the *ARNTL* gene) and CLOCK expression (Crumbley & Burris, 2011; Guillaumond et al., 2005). Over the past few years, small molecule agonists and antagonists have been developed or discovered that target many of these factors (Chen et al., 2018; Ribeiro et al., 2021), thus raising the possibility that the clock and clock-regulated processes can be modulated pharmacologically.

Based on our previous report that treatment of U2OS osteosarcoma cells with a combination of the CRY inhibitor KS15 and REV-ERB antagonist SR8278 was associated with increased expression of XPA and WEE1 (Anabtawi et al., 2021), we were curious whether the low levels of XPA at noon in human skin could be increased by treatment with these compounds. We therefore topically treated human skin explants obtained from routine panniculectomies with a combination of KS15 and SR8278 (100 µM each) or with DMSO vehicle at 5-6 pm and then examined the epidermal expression of the clock-controlled genes *XPA, WEE1*, and *CDKN1A* and four core clock genes (*ARNTL, NR1D2, CRY1*, and *PER2*) 18 hours later by RT-qPCR. The concentrations of these compounds were selected based on the expected higher concentration needed to penetrate the stratum corneum than concentrations typically used in cell culture in vitro. The fold-change in expression in the KS15+SR8278 (KS+SR)-treated samples relative to DMSO-treated skin samples is provided in **Figure 1B** along with the age and sex of each skin donor.

Though there was significant inter-individual variation in responses to the treatment, we noted that expression levels of both *XPA* and *WEE1* were significantly elevated by the treatment with KS15 and SR8278 (**Figure 1C**; lower ΔC_t_ indicates higher expression), with 7-8 of the 16 skin samples exhibiting at least a two-fold increase in mRNA levels of these two gene products. Analysis of the core circadian genes revealed that only *PER2* displayed a trend towards increased expression following treatment with the cryptochoursome and REV-ERB inhibitors (**Figure 1C**). This could be due in part to the greater variability in circadian gene expression and times of peak expression that exist in human skin explants (Cvammen & Kemp, 2024a).

Given that higher expression of *XPA* and *WEE1* may promote nucleotide excision repair and enhanced G2/M cell cycle arrest to limit the negative effects of UV-induced DNA damage, we exposed a subset of these skin biopsies to UVB radiation at 12 pm (18 hours after treatment with DMSO or KS15 and SR8278) and then fixed the tissue in formalin 24 hours later for analysis of apoptotic, “sunburn” cells by H&E staining (Sheehan & Young, 2002). As shown in **Figure 1D**, the fold-change in number of apoptotic cells was found to be reduced by KS15 and SR8278 treatment in each of the five samples.

Though this work suggests the possibility that the skin circadian clock can be modulated pharmacologically to alter responses to UV radiation and potentially other stressors, we note that the pilot study described here has several limitations, including a relatively small sample size and the use of skin explants from donors of a wide range of ages (age 17-67) of unknown health status. Moreover, we used only a single concentration of KS15 and SR8278 and monitored changes in gene expression at only the mRNA level (not protein level) of a very limited number of genes at only a single time point. Finally, we recently showed that both KS15 and SR8278 absorb within the UVB spectrum (Cvammen & Kemp, 2024b) and that SR8278 has off-target effects (Ushaswini et al., 2026), and thus the apparent photoprotection observed here could be due in part to the direct absorption of UVB light and not solely due to changes in clock gene function or to off-target effects of the drugs.

Nonetheless, to our knowledge, this is the first study to make use of circadian clock-modulating drugs in the context of human skin. As additional natural products and more portent small molecules that target the circadian clock continue to be discovered and characterized (Maram et al., 2026), our data here argue that there may be value in exploring their use on human skin and in UV photoprotection.

## Methods

The transcriptomic data set GSE205155 (Del Olmo et al., 2022) was downloaded from the Gene Expression Omnibus database. Raw data was extracted and normalized with the limma R package and then normalized to the highest expression time point for each individual subject. Human skin from obtained from routine panniculectomies at Miami Valley Hospital. Patients provided written, informed consent, and the study was provided by the Wright State University Institutional Review Board. Other than the age and sex of the patients, no other identifying information about the patients was obtained. The skin donations were collected from at variable times between the hours of 10:30 AM and 4:30 PM within approximately 30 minutes following the surgery. Skin samples were washed with tap water to remove excess blood, and then epidermis was briefly washed with 70% ethanol. Small, 8 mm punch biopsies were placed in Millicell cell culture inserts in wells of a 24-well plate containing 100 µl of basal DMEM containing penicillin/streptomycin as previously described (Kemp et al., 2019) and then incubated in a 5% CO_2_ 37°C incubator. At 5-6 pm, 25 µl of either DMSO or a combination of 100 mM KS15 and SR8278 (in DMSO) were aliquoted onto the top of the biopsy. The skin samples were returned to the incubator until 11-12 pm the next day. Biopsies were then stored in RNA*Later* at −20°C. RNA was purified from the epidermis following heat shock separation from the dermis (Cvammen & Kemp, 2024a) and then reverse transcribed with a QuantiTect Reverse Transcription Kit (Qiagen). cDNA was amplified using 2X TaqMan Fast Universal PCR Master Mix and TaqMan probes on an Azure Cielo 6 real-time PCR instrument. Beta-2 microglobulin (B2M) was used as the housekeeping gene for normalization purposes between different samples. A small number of these skin biopsies were also exposed to 1,200 J/m^2^ UVB radiation using UVP XX-15M UVB bulbs (Analytik-Jena). Biopsies were fixed in formalin, sectioned, and stained with hematoxylin and eosin to identify and count apoptotic cells.

## Acknowledgments

We thank the WSU Proteome Analysis Laboratory for the use of equipment to carry out this work.

## Author Contributions

William Cvammen: Data curation, Writing – review and editing Michael G. Kemp: Conceptualization, Data curation, Writing – original draft, Funding acquisition, Project administration

